# Are sexually selected traits affected by a poor environment early in life?

**DOI:** 10.1101/045740

**Authors:** Regina Vega-Trejo, Michael D. Jennions, Megan L. Head

## Abstract

Conditions experienced early in life can affect key life-history traits. Individuals that experience a poor nutritional environment early in life can reduce potential costs by delaying sexual maturation. The direct costs of delaying maturation are well known (i.e. delayed onset of breeding), but individuals can also face additional costs as adults. Some of these ‘hidden costs’ accrue due to cryptic morphological and physiological changes. In mosquitofish (*Gambusia holbrooki*), males with limited food intake early in life delay maturation to eventually reach a similar adult body size to their well-fed counterparts (‘catch-up growth’). Here we tested whether a poor diet early in life incurs hidden costs due to reduced expression of sexually selected male traits (genital size and ejaculate traits). We found that diet early in life significantly influenced sperm reserves and replenishment rate due to age and development-rate dependent effects. Although catching-up in body size ameliorates some of the costs of a poor start in life for males, our findings suggest that long-term fitness costs might arise because of sexually selection against these males. Our results highlight that fitness costs of a poor start in life can extend into adulthood.

## Introduction

Conditions experienced early in life can affect the life-history trajectories of individuals (Monaghan, 2008). In particular, limited food availability during early development can lower growth rates and, ultimately, reduce adult body size (Dmitriew, 2011). Given the consequences of a poor nutritional environment, organisms often respond in ways that reduce potential fitness costs of smaller body size. For example, when food again becomes available, individuals can compensate for a poor start in life by either accelerating their growth (compensatory growth) or by taking longer to reach maturity, eventually attaining a similar adult size to well-fed individuals (catch-up growth; reviews: Metcalfe & Monaghan, 2001; Hector & Nakagawa, 2012). Although there are usually clear benefits to attaining large adult size, such as increased survival and improved reproductive success (e.g. Lee *et al.,* 2013), there can be costs (e.g. Yearsley *et al*., 2004). For instance, an obvious cost of catch-up growth is delayed maturation, which can increase generation time and decrease reproductive lifespan (Roff, 1992; Marcil-Ferland *et al*., 2013). Delayed maturation and other immediate costs of growth plasticity are well documented (Oli *et al*., 2002; Ali *et al*., 2003; Hector & Nakagawa, 2012; Teder *et al*., 2014). However, less is known about delayed costs that arise because of poor conditions early in life, and how they affect allocation to different life history traits (but see: Barry, 2013; Runagall-McNaull *et al*., 2015).

Poor nutrition early in life can cause changes in adult behaviour (Ali *et al*., 2003; Royle *et al*., 2005), morphology (Ohlsson & Smith, 2001; Chin *et al*., 2013), and key life-history traits (Lindstrom *et al*., 2005; Taborsky, 2006; Mugabo *et al*., 2010; Orizaola *et al*., 2014). These changes often impose a cost, such as reduced locomotor performance (Hector *et al*., 2012), or reduced sexual ornament expression (Ohlsson *et al*., 2002). However, the effect of a poor start in life on the expression of some traits is less obvious. Even if there is no effect on adult external morphology, fitness can still decrease. This is often referred to as a hidden cost. For example, adults on a low quality diet can be indistinguishable morphologically from those on a standard diet, but suffer a reduction in hidden traits such as telomere length or plasma antioxidant levels, which reduce lifespan (Birkhead *et al*., 1999; Blount *et al*., 2003; Reichert *et al*., 2015). A poor diet early in life is generally assumed to reduce adult fitness. While most investigations of hidden costs focus on naturally selected traits, costs associated with the expression of sexually selected traits, which are equally important have received far less attention (but see: Zajitschek *et al*., 2009; Sentinella *et al*., 2013; Fricke *et al*., 2015).

Most sexually selected traits are under strong directional selection, and developmental nutrition is often a major determinant of their expression. Pre-copulatory traits (i.e. sexually selected traits that affect mating success) can be detrimentally affected by poor early nutrition. For example, eye-span in stalked-eyed flies and song repertoire size in great reed warblers, which are both traits that affect female mate choice, are negatively affected by nutritional stress during development (David *et al*., 2000; Nowicki *et al*., 2000). This reduction in investment could reflect a trade-off between sexually and naturally selected traits. For example, when early nutritional conditions are poor, males sometimes reduce investment in sexual ornaments to maintain their oxidant defence systems (Ohlsson *et al*., 2002; Blount *et al*., 2003). Similar trade-offs could also affect investment into different sexually selected traits. In general reproductive success depends on both pre‐ and postcopulatory fitness components (Andersson, 1994; Fedina & Lewis, 2008; Tigreros, 2013). Given nutritional stress and lower resource availability, individuals might invest differently in precopulatory and post-copulatory traits. This could result in a shift in the relative allocation of resources to post-copulatory competitiveness versus attractive ornaments (Cordes *et al*., 2015). For example, greater investment in larger body size or weaponry can result in smaller testes (Droney, 1998; Somjee *et al*., 2015).

Fertilization success in males is highly dependent on resource allocation to traits that are under postcopulatory sexual selection, especially in species with high levels of sperm competition (Parker & Pizzari, 2010). In such species, males tend to invest in relatively larger testes that produce more sperm (Wedell *et al*., 2002). However, the amount of sperm a male can produce is not the sole predictor of his sperm competitiveness. Sperm competition outcome is also dependent on sperm viability, swimming speed, and, in some cases, length (Boschetto *et al*., 2011; Mautz *et al*., 2013). Since ejaculates are expected to be costly to produce (Olsson *et al*., 1997; Kahrl & Cox, 2015), it follows that a poor diet could negatively impact the number, quality, and rate of sperm production (e.g. O'Dea *et al*., 2014; Bunning *et al*., 2015). Although condition dependence of sperm traits has been examined in several species, this is usually only in the context of the short-term effect of a recently manipulated adult diet (e.g. Lewis & Wedell, 2007; Gasparini *et al*., 2013; Kahrl & Cox, 2015). It is rarely tested whether a restricted diet early in life, followed by a return to a regular diet, affects sperm traits (but see: Tigreros, 2013; Cordes *et al*., 2015; Rosenthal & Hebets, 2015).

In this study, we test whether the diet of juvenile male mosquitofish *(Gambusia holbrooki)* affects adult sexual traits. Previous studies have shown that males with limited food intake early in life reach a similar size to males on a normal diet, albeit with a delay in maturation (i.e. catch-up growth; Livingston et al. 2014; R. Vega-Trejo, M.L. Head, and M.D. Jennions ‘unpublished data’). Additionally, it has been shown that males with a poor start in life are less attractive to females than those that develop normally (Kahn *et al*., 2012). Mosquitofish are poecillid fishes characterised by a coercive mating system, frequent mating attempts, and intense sexual selection, including sperm competition (Bisazza & Marin, 1991; 1995; O’Dea *et al*., 2014). Mosquitofish exhibit internal fertilization in which males inseminate females using an intromittent organ modified from the anal fin called gonopodium (Pyke, 2005). Here we test whether males initially raised on a restricted diet incur hidden costs due to reduced expression of traits that are important in post-copulatory sexual selection, specifically due to the production of lower quality ejaculates.

## Methods

Our data are from fish that were bred as part of a larger study testing how inbreeding and food restriction affect compensatory growth (R. Vega-Trejo, M.L. Head, and M.D. Jennions ‘unpublished data’). In that study we found no effects of inbreeding on any of the life history variables measured (growth trajectories and adult size). Here, we are specifically interested in whether diet restriction during early life influences ejaculate traits so, for clarity, we analyse the data without including inbreeding in the models. Including inbreeding and its interactions does not qualitatively alter our results because these terms all had very small, and non-significant effects (see Supporting information 1; Note to reviewers: a subset of this data will be published in J. Marsh, R. Vega-Trejo, M.L. Head, and M.D. Jennions ‘unpublished data’; which is a study explicitly linking inbreeding to sperm production and to share of paternity.).

We used mosquitofish descended from fish captured from ponds in Canberra, Australia. In each experimental block we mated individuals from two families (e.g. A and B in block 1, C and D in block 2 and so on). Brothers and sisters from full sibling families were paired to create inbred offspring (AA, BB) and outbred offspring with reciprocal male-female crosses (AB, BA) to generate four cross-types. We set up 29 blocks. Offspring from these pairings were reared individually in 1L tanks until maturity. They underwent a diet manipulation for 21 days between day 7 and day 28-post birth. Fish on the control diet were fed *ad libitum* with *Artemia nauplii* twice a day (i.e. our standard laboratory feeding regime) whereas fish on the restricted diet were fed 3mg of wet mass adult *A. nauplii* once every other day (for justification see Livingston et al. 2014). Broods were split between the control and restricted diet treatment. This experimental design allowed us to investigate how early diet influences male sexual traits after maturity. Males were considered mature when their gonopodium was translucent with a spine visible at the tip (Stearns, 1983; Zulian *et al*., 1993). We collected body size and sperm data from mature males (range: 2 to 18 weeks post-maturity). We define ‘developmental time’ as the number of days that males took to reach maturity, and ‘adult age’ as the age at which sperm was extracted (i.e. total age – developmental time).

### SPERM TRAITS

We tested 452 males from 192 broods. Sperm was collected on three occasions: on day 0 we stripped virgin males of sperm (see below) to obtain a measure of their maximum sperm reserves, one day later we stripped males to obtain a measure of their sperm replenishment rate (i.e. the amount of sperm produced in a 24 hr period), on day 3 we stripped males to measure sperm velocity. We also calculated the proportion of sperm replenished (= number of sperm at day 1 / number of sperm at day 0). We arcsin-transformed the proportion to normalize the error distribution.

### SPERM COLLECTION

To strip ejaculates, males were anaesthetized in ice-cold water. Each male was then placed on a glass slide previously coated with 1% PVA (this coating prevents sperm bundles sticking to the slide and aids their collection) under a dissecting microscope and his gonopodium was swung forward. We applied gentle pressure to the abdomen to eject all the available sperm. We transferred the ejaculate to an Eppendorf tube with 100 - 900 μL of extender medium (207 mM NaCl, 5.4 mM KCl, 1.3 mM CaCl_2_, 0.49 mM MgCl_2_, 0.41 mM MgSO_4_, 10 mM Tris, pH 7.5) depending on the amount of ejaculate stripped. Sperm remains quiescent in this solution (Gardiner, 1978). After this procedure males were returned to their individual tanks. Sperm collection was done blind to male treatment and only performed by RVT.

### SPERM NUMBER

To estimate the number of sperm we vortexed the sperm solution for one minute and then mixed it repeatedly with a pipette (between 20 and 30 times) to break sperm bundles and distribute the sperm evenly throughout the sample. We placed 3 μL of the solution on a 20 micron capillary slide (Leja) and counted the sperm using CEROS Sperm Tracker under a 100× magnification. We counted five subsamples per sample. Repeatability was estimated following Nakagawa and Schielzeth (2010) using the *rptR* package in *R 3.0.2* (R Development Core Team, 2012). Repeatability within samples was very high for sperm numbers (sperm at day 0: *r* = 0.85 ± 0.01 SE; sperm at day 1: *r* = 0.91 ± 0.006 SE). The mean of the five subsamples was used for further analyses. The threshold values defining cell detection were predetermined as elongation percentage 15-65, head size 5 - 15 μm, and the static tail filter was set off. Sperm were counted blind to male treatment.

### SPERM VELOCITY

For each ejaculate we analysed three samples. For each sample we collected 3 μL of the diluted sperm (above) and placed this in the centre of a cell of a 12- cell multitest slide (MP Biomedicals, Aurora, OH, USA) previously coated with 1% polyvinyl alcohol solution to avoid sperm sticking to the slide. The sample was then activated with a 3 μL solution of 150 mM KCl and 2 mg ml^-1^ bovine serum albumin (Billard & Cosson, 1992) and covered with a cover slip. We analysed sperm velocity within 30 seconds of activation for three subsamples to increase the number of sperm contributing to our mean velocity measure. We used an average of 109.3 ± 3.49 SE sperm tracks per ejaculate (minimum 10 sperm tracks / male) to estimate sperm velocity. We excluded six males from the velocity analysis because they had fewer than 10 sperm tracks. We recorded two standard measures of sperm velocity: (1) average path velocity (VAP), which estimates the average velocity of sperm cells over a smoothed cell path and (2) curvilinear velocity (VCL), the actual velocity along the trajectory using CEROS Sperm Tracker (Hamilton Thorne Research, Beverly, MA, USA). The threshold values defining static cells were predetermined at 20 μm/s for VAP and 15 μm/s for VCL. Repeatability within samples was high for the parameters (VAP: *r* = 0.65 ± 0.02 SE; VCL: *r* = 0.58 ± 0.03) and so the mean of the three subsamples was used in our analyses. Due to the near perfect correlation between VAP and VCL (*r*=0.961, P<0.001) we only use VAP in our analyses.

### MALE MORPHOLOGY

All males were measured a week after the sperm extractions. Males were anaesthetized by submersion in ice-cold water for a few seconds to reduce movement and then placed on polystyrene with a microscopic ruler (0.1 mm gradation) to be photographed. We measured male standard length (SL = snout tip to base of caudal fin) and gonopodium length using Image J software (Abramoff *et al*., 2004).

### STATISTICAL ANALYSIS

We removed one male (of 452) from the analysis because he had a higher number of sperm on day 1 than day 0 indicating that not all sperm were collected during the first extraction.

To analyse the effect of diet treatment on male sexual traits we used generalized linear mixed models (GLMM). We constructed separate models for each of our five response variables: gonopodium length, number of sperm at day 0, number of sperm at day 1 (i.e. replenishment rate), proportion of sperm replenished (arc-sine transformed), and VAP. In each model we included diet as a fixed factor, and male standard length, development time, and adult age as fixed covariates, as well as including all two-way interactions with diet. Both gonopodium length and body size were logtransformed. Adult age was not included in the model for gonopodium length as there is almost no post-maturity growth in *G. holbrooki*.

There were significant bivariate correlations between development time and body size (*r*= 0.62, P<0.001) as well as development time and adult age (*r* = −0.77, P<0.001). The former reflects a biological relationship and the latter is due to a logistic constraint (i.e. having to terminate the experiment). Even so, these correlations were not so large as to preclude including all three terms as covariates in a GLMM due to colinearity problems (running each model with only one covariate at a time produce comparable effect sizes for focal terms).

More importantly, we needed to take into account that mean development time and mean adult age differed significantly between the diets (separate GLMMs with diet as the single fixed factor, and random factors as below: both *P* <0.001). Including development time and/or adult age as raw covariates could obscure a main effect of diet (i.e. these are covariates measured post-treatment *sensu* Gelman and Hill (2007), p.188 that might causally mediate any diet effect). We therefore standardised each covariate (mean=0, S.D =1) *within* each diet treatment. Although male standard length did not differ between diet treatments (*P* = 0.451; see Results), we standardised it within each diet for consistency.

Centering the covariates within each diet affects their interpretation. The effect of adult age and development time (or either terms interaction with diet) should be interpreted as age or development rate relative to other males on the same diet. The main effects of diet are then interpretable at those for a male of average age, size, and development time for its treatment type (see Schielzeth, 2010). We should note, however, that despite concerns about including posttreatment covariates, the standardization of covariates across the full data set, rather than within diet treatments, does not affect our main conclusions (Supporting information 2).

In all the GLMMs we specified a Gaussian error distribution and checked the distribution of model residuals to ensure this was appropriate. The use of Poisson error (for count data) and binomial error (for proportions) provided a worse fit to the data than the use of Gaussian error on the raw or transformed dependent variables. Each model was fitted using the *lme4* package in *R 3.0.2* software with block, maternal identity, and sire identity as random factors (see Vega-Trejo *et al*., 2015). All model terms were tested for significance using the Anova function in the *car* package specifying Type III Wald chi-square tests. Model simplification (i.e. removing non-significant interaction and main terms) did not change our results. Figures are presented using raw data rather than model predictions (i.e. ignoring random effects). Summary statistics are presented as mean ± SE.

## Results

The correlations between the four ejaculate traits are provided in Table 1. Diet treatment means for all traits are provided in Table 2. Parameter estimates are provided in Table 3.

**Table 1.** Correlations between sperm traits measured. Estimates are followed by p-values in brackets. N = 452 males

| | Number of sperm at day 1 | Proportion of sperm replenished | Sperm velocity (VAP, $\mu\text{m/s}$) |
| --- | --- | --- | --- |
| Number of sperm at day 0 | 0.468 (<0.001) | -0.117 (0.015) | 0.055 (0.271) |
| Number of sperm at day 1 |  | 0.664 (<0.001) | 0.032 (0.526) |
| Proportion of sperm replenished |  |  | 0.059 (0.246) |

**Table 2.** Treatment means ± SE (n) for the five traits measured.

|  | Control diet | Restricted diet |
| --- | --- | --- |
| Gonopodium length (mm) | 6.94 ± 0.05 (223) | 7.03 ± 0.04 (226) |
| Number of sperm at day 0 ($\times 10^5$) | 194.58 ± 6.37 (225) | 176.20 ± 6.14 (227) |
| Number of sperm at day 1 ($\times 10^5$) | 62.36 ± 2.72 (225) | 47.92 ± 2.67 (227) |
| Proportion of sperm replenished | 0.35 ± 0.02 (225) | 0.29 ± 0.01 (227) |
| Sperm velocity (VAP, $\mu\text{m/s}$) | 83.10 ± 1.11 (225) | 81.88 ± 1.12 (227) |

**Table 3.** Results from mixed models with parameter estimates and chi square (χ^2^) tests for food treatment, size, developmental time, and adult age. P-values in bold indicate significant values. Covariates were standardised within food treatment. The sample sizes for control and restricted diets are 225 and 227, respectively.

| | Predictor | Estimate | SE | $\chi^2$ | P |
| --- | --- | --- | --- | --- | --- |
| Gonopodium length [ln(mm)] | Intercept | 0.842 | 0.002 | $1.553 \times 10^5$ | **<0.001** |
|  | Diet (control) | -0.004 | 0.001 | 5.935 | **<0.001** |
|  | Size | 0.031 | 0.002 | 138.02 | **<0.001** |
|  | Developmental time | 0.006 | 0.002 | 12.249 | **<0.001** |
|  | Diet × Size | 0.010 | 0.002 | 10.530 | **0.001** |
|  | Diet × Developmental time | -0.003 | 0.001 | 2.214 | 0.074 |
| Number of sperm at day 0 (total count) | Intercept | $185.293 \times 10^5$ | $7.010 \times 10^5$ | 698.634 | **<0.001** |
| | Diet (control) | $7.802 \times 10^5$ | $4.000 \times 10^5$ | 3.804 | 0.051 |
| | Size | $25.848 \times 10^5$ | $5.544 \times 10^5$ | 21.738 | **<0.001** |
| | Adult age | $9.754 \times 10^5$ | $7.042 \times 10^5$ | 1.918 | 0.166 |
| | Developmental time | $-15.438 \times 10^5$ | $7.899 \times 10^5$ | 3.820 | 0.051 |
| | Diet × Size | $2.644 \times 10^5$ | $5.573 \times 10^5$ | 0.225 | 0.635 |
| | Diet × Adult age | $-26.279 \times 10^5$ | $6.355 \times 10^5$ | 17.099 | **<0.001** |
| | Diet × Developmental time | $-11.169 \times 10^5$ | $7.415 \times 10^5$ | 2.269 | 0.132 |
| Number of sperm at day 1 (total count) | Intercept | $54.973 \times 10^5$ | $2.410 \times 10^5$ | 520.288 | **<0.001** |
| | Diet (control) | $6.449 \times 10^5$ | $1.753 \times 10^5$ | 13.529 | **<0.001** |
| | Size | $13.269 \times 10^5$ | $2.534 \times 10^5$ | 27.400 | **<0.001** |
| | Adult age | $2.651 \times 10^5$ | $2.973 \times 10^5$ | 0.795 | 0.372 |
| | Developmental time | $-16.806 \times 10^5$ | $3.471 \times 10^5$ | 23.435 | **<0.001** |
| | Diet × Size | $1.054 \times 10^5$ | $2.552 \times 10^5$ | 0.170 | 0.679 |
| | Diet × Adult age | $-7.211 \times 10^5$ | $2.776 \times 10^5$ | 6.747 | **0.009** |
| | Diet × Developmental time | $-4.724 \times 10^5$ | $3.345 \times 10^5$ | 1.999 | 0.157 |
| Proportion of sperm replenished | Intercept | 202.833 | 6.269 | 1046.915 | **<0.001** |
|  | Diet (control) | 16.163 | 5.661 | 8.153 | **0.004** |
|  | Size | 24.665 | 8.022 | 9.453 | **0.002** |
|  | Adult age | 0.683 | 9.086 | 0.006 | 0.940 |
|  | Developmental time | -51.155 | 10.757 | 22.615 | **<0.001** |
|  | Diet × Size | -0.279 | 8.045 | 0.001 | 0.972 |
|  | Diet × Adult age | -4.323 | 8.884 | 0.237 | 0.626 |
|  | Diet × Developmental time | -4.646 | 10.626 | 0.191 | 0.662 |
| Sperm velocity (VAP, $\mu\text{m/s}$) | Intercept | 82.922 | 0.956 | 7524.697 | <0.001 |
|  | Diet (control) | 0.500 | 0.756 | 0.438 | 0.508 |
|  | Size | 0.379 | 1.101 | 0.118 | 0.731 |
|  | Adult age | -6.372 | 1.300 | 24.022 | <0.001 |
|  | Developmental time | -2.078 | 1.509 | 1.894 | 0.169 |
|  | Diet × Size | -0.719 | 1.121 | 0.412 | 0.521 |
|  | Diet × Adult age | -0.489 | 1.209 | 0.164 | 0.686 |
|  | Diet × Developmental time | 1.784 | 1.455 | 1.505 | 0.219 |

### EFFECT OF TREATMENT ON MALE MORPHOLOGY

There was no significant difference between the diet treatments in male body size at maturity (control: 23.51 ± 0.14 mm; restricted diet: 23.35 ± 0.11 mm). Against expectations (see Livingston et al. 2014), males on the restricted diet generally had a *larger* gonopodium than control diet males for males that were of average or smaller body size (diet by size interaction, *P* = 0.001; Fig.1). Correcting for size, males that took relatively longer to mature on a given diet had a significantly longer gonopodium (P < 0.001). There was no significant interaction between development time and diet (P = 0.074).

### WHAT INFLUENCES SPERM TRAITS?

#### Initial sperm reserves (Day 0)

As expected, larger males had significantly greater initial sperm reserves (P < 0.001), but this relationship did not differ between the two diets (P = 0.635). However, diet significantly influenced the extent to which sperm reserves increased with age (diet by age interaction, P < 0.001). Initial sperm reserves of males on the restricted diet increased significantly with age, while there was no such change with age for males on the control diet (i.e. parameter estimate for age is for control males, P = 0.166; Fig.2). There was a marginally non-significant effect of development time on initial sperm reserves (P = 0.051).

#### Sperm replenishment rate (Day 1)

Larger males had significantly higher sperm replenishment rates than smaller males (P < 0.001); but, controlling for body size, males that took longer to develop had a lower sperm replenishment rate (P < 0.001). For an individual of average age, size and development time for its diet treatment, a male reared on the control diet had a significantly higher replenishment rate (P < 0.001). There was, however, a significant difference between the diets in how age was related to replenishment rate: males on the restricted diet had a significantly higher replenishment rate when older (age by diet interaction, P = 0.009), but there was no age-dependent change for control males (P = 0.372).

Overall, males on the control diet replenished a greater proportion of their initial sperm reserves within 24 h (P = 0.004). The proportion replenished was significantly greater for larger males (P = 0.002) and for males with a shorter development time (P < 0.001). There was no significant effect of male age (P = 0.940), nor were interactions between diet and size or age or development time significant (all P > 0.62).

Finally, there was no effect of diet on sperm velocity (VAP). Sperm velocity did, however, decrease significantly with adult age (P < 0.001; Fig.3), but it was unrelated to development time or male size. There were no significant interactions between diet and size or age or development time (all P > 0.22).

## Discussion

Phenotypic responses to nutritional constraints early in life can affect an individual’s fitness due to changes in their adult performance. Like many species (Metcalfe & Monaghan, 2001; Barry, 2013), juvenile mosquitofish experiencing a poor diet early in life extend their development time to attain a similar body size to individuals on good diets (see Livingston et al. 2014; R. Vega-Trejo, M.L. Head, and M.D. Jennions ‘unpublished data’). Here we tested whether a poor diet early in life has additional “hidden costs” for males because it reduces ejaculate quality. We found that early diet had a significant influence on initial sperm reserves, sperm replenishment rate, and the proportion of sperm replenished, but not on sperm velocity. Males that had their diet restricted during development had smaller sperm reserves and lower replenishment rates early in adulthood than males that did not have a restricted diet. However, males on both diets attained similar sperm reserves and replenishment rates when older, as indicated by an interaction between diet and adult age. Our results combined with those from our previous studies (Kahn *et al*., 2012; Livingston *et al*., 2014) suggest that a poor diet early in life can have multiple costs, not only delayed maturation, but also due to additional ‘hidden’ costs experienced as adults.

Sperm production has been shown to be condition dependent in a variety of species: when the diet of adults is restricted they have smaller sperm reserves (e.g. Perry & Rowe, 2010; Rahman *et al*., 2014; Kahrl & Cox, 2015) and lower replenishment rates (e.g. O'Dea *et al*., 2014). There are far fewer studies that explore the effects of a poor diet only experienced early in life on sperm reserves and replenishment rates (but see: Tigreros, 2013; Cordes *et al*., 2015). We found that both sperm reserves and replenishment rates were affected by a male’s early diet in an age-dependent manner. Thus, in addition to the immediate condition-dependence of sperm production reported in previous studies, we have shown that early diet restriction can have delayed, long-term effects. Sperm number is often a good predictor of male fertilization success under sperm competition (Parker, 1990), so such dietary effects are likely to have important fitness consequences. Our results suggest that even if males that had a poor start in life mate as often as those that had a regular diet, their lower sperm reserves and reduced ability to continuously produce ejaculates could decrease their reproductive success. Although sperm reserves increase with time after sexual maturity (here and Evans *et al*., 2002), males experiencing restricted diets during development would still be disadvantaged because they take longer to reach full sperm reserves.

The quantity and quality of food acquired during early life stages can generate different optimal adult reproductive strategies (Cordes *et al*., 2015). Our results show that early diet has important effects on traits that influence reproductive success. Our findings are comparable to those in studies showing that the conditions experienced early in life due to differences in parental care can affect adult traits that are important for reproductive success (Hunt & Simmons, 2000; Reinhold, 2002). Parental care itself can improve offspring quality leading to an increase in offspring reproductive success (Klug *et al*., 2012). In addition, the diet or conditions that parents experience can be transmitted to their developing offspring (i.e., transgenerational effects: (Monaghan, 2008; Alonso-Alvarez & Velando, 2012; Klug & Bonsall, 2014). For example, in a dung beetle, developing larvae depend on nutrients provided by their parents which affects offspring body and horn size and ultimately their reproductive success (Hunt & Simmons, 2000). Similarly, in birds the amount of carotenoids available to mothers can influence the amount of these pigments that is deposited into egg yolks, which can then affect the ornamental coloration offspring display as adults (McGraw *et al*., 2005). Our results highlight that regardless of whether early nutritional environment is determined by parental care, an individual’s own resource acquisition ability, or the habitat into which an animal is born, the effects on fitness can be far reaching, extending well into adulthood.

We found that sperm velocity decreases with adult age. Sperm quality is expected to decline with age due to a reduction in its fertilising efficiency and/or the genetic quality of sperm produced by ageing males (Pizzari *et al*., 2008). This expectation is supported by empirical studies showing that sperm velocity deteriorates with male age (e.g. Sloter *et al*., 2006; Moller *et al*., 2009; Cornwallis *et al*., 2014). The poor sperm quality of older males has been shown to lower fertilization success under sperm competition (i.e. when competing with sperm from younger males; e.g. Radwan *et al*., 2005) but not others (Hoysak *et al*., 2004; Gasparini *et al*., 2010). There are two possible reasons why older males might have lower quality sperm. One is that old males *produce* lower quality sperm (an effect of male age per se e.g. Jones *et al*., 2007; Moller *et al*., 2009). The other is that the sperm of older males has deteriorated due to spending more time in storage (an effect of sperm age e.g. Siva-Jothy, 2000; Reinhardt & Siva-Jothy, 2005). In our study all sperm velocity measures were obtained from sperm that were a maximum of three days old, so the observed lower sperm velocity is due to an effect of male age rather than sperm age. It is intriguing that sperm numbers increased with age (at least for males on the low food diet), while sperm velocity declined with age for all males. This suggests that sperm number might be a more important determinant of fertilization success than sperm velocity, and is therefore more likely to be maintained given limited resources. This is supported by data from other Poeciliids showing that sperm number is more important in postcopulatory sexual selection than sperm velocity (see Gasparini *et al*., 2013). Interestingly, despite the effect of adult age on sperm velocity, unlike the case for sperm numbers, there was no interaction between male age and diet.

Male fitness depends on the ability to acquire mates and then to gain paternity when females mate multiply. In mosquitofish, the quantity and the quality of sperm are not the only traits that influence reproductive success. We also measured a morphological trait, gonopodium size, which can affect female mate choice in Poeciliids (Brooks & Caithness, 1995; Langerhans *et al*., 2005; Kahn *et al*., 2010; Devigili *et al*., 2015) and has been implicated as a potentially important trait affecting sperm transfer (Pyke, 2005; Head *et al*., 2015). We found that small and medium-sized males on a restricted diet early in life had longer gonopodia than those on a regular diet (but see Livingston et.al. 2014 for a different finding). However, this effect is unlikely to lead to important fitness consequences because the dietary effect was only evident for small to medium sized males. A female preference for males with a longer gonopodium has been shown in *G. holbrooki,* but it is only evident for large bodied males (Kahn *et al*., 2010). In addition, insemination success seems to depend on both male body size and gonopodium length - males with a relatively longer gonopodium are likely to be more successful only when they are large (Head *et al*., 2015). Finally, due to the coercive behaviour on mosquitofish, males with larger gonopodia could have a higher reproductive success if males with a larger gonopoidum would be more successful at harassing females by performing more gonopodial thrusts (Elgee 2012, Evans 2011, Reynolds 1993) or their ability to perform sneak copulations (Pilastro 1997)”. Hence, the effects of diet on relative gonopodium length we observed are, based on current evidence, unlikely to affect male reproductive success.

In sum, some ejaculate traits in *G. holbrooki* depend on the diet experienced early in life and male age. The observed interaction between early diet and adult age suggests a hidden cost via effects on sperm reserves and replenishment rates. A previous study showed that early life diet influences male attractiveness in *G. holbrooki* (Kahn *et al*., 2012). Together these studies suggest that early diet could have fitness consequences well into adulthood. Our findings are similar to those in other species where males on different diets superficially look the same, but differ in social dominance (e.g. Royle *et al*., 2005), telomere length or plasma antioxidant levels (Blount et.al. 2003, Reichert et.al. 2015). As with these studies it is assumed that the affected traits will alter male fitness, but the actual effects on male fitness of a poor early diet due to changes in sperm traits, genital size or other life history traits remain unclear. Ideally, future studies should directly measure the relative reproductive success of males that undergo a poor start in life in a competitive mating context.

## Acknowledgments

We thank the ANU Animal Services team for fish maintenance. This work was supported by the Australian Research Council (DP120100339). Animal use permit: ANU AEEC animal ethics protocol A2011/64. R.V.-T. is supported by fellowships from Consejo Nacional de Ciencia y Tecnoloífa-México and the Research School of Biology.

---

Figure 1. Effect of diet treatment on the relationship between male body size and gonopodium length. Standard length is standardised within each diet. Black symbols and lines represent the control diet, grey symbols and lines represent the restricted diet. Lines represent simple regressions ignoring random effects. Grey shading represents 95% confidence intervals.

Figure 2. Effect of diet treatment on the relationship between number of sperm at day 0 and adult age. Number of sperm represent total counts, adult age is standardised within each diet. Black symbols and lines represent the control diet, grey symbols and lines represent the restricted diet. Lines represent simple regressions ignoring random effects. Grey shading represents 95% confidence intervals.

Figure 3. The relationship between VAP (average path velocity) and adult age. VAP is given in μm/s, adult age is standardised within each diet. The line represents a simple regression ignoring random effects. Grey shading represents 95% confidence intervals.

